# Crl activates transcription by stabilizing the active conformation of the master stress factor σ^S^

**DOI:** 10.1101/721944

**Authors:** Juncao Xu, Kaijie Cui, Liqiang Shen, Jing Shi, Lingting Li, Linlin You, Chengli Fang, Guoping Zhao, Yu Feng, Bei Yang, Yu Zhang

**Author notes:** These authors contributed equally to this work (Juncao Xu, Kaijie Cui). Corresponding authors (Y. Z.); (Y. F.); (G. Z.); (B. Y.). Lead Contact (Y. Z.).

## Abstract

σ^S^ is a master transcription initiation factor that protects bacterial cells from various harmful environmental stresses and antibiotic pressure. Although its mechanism remains unclear, it is known that full activation of σ^S^-mediated transcription requires a σ^S^-specific activator, Crl. In this study, we determined a 3.80 Å cryo-EM structure of an *E. coli* transcription activation complex (*E. coli* Crl-TAC) comprising *E. coli* σ^S^-RNAP holoenzyme, Crl, and a nucleic-acid scaffold. The structure reveals that Crl interacts with the domain 2 of σ^S^ (σ^S^_2_), sharing no interaction with promoter DNA. Subsequent hydrogen-deuterium exchange mass spectrometry (HDX-MS) results indicate that Crl stabilizes key structural motifs of σ^S^_2_ to promote the assembly of σ^S^-RNAP holoenzyme and also to facilitate formation of the RNA polymerase-promoter DNA open complex (RPo). Our study demonstrates a unique DNA contact-independent mechanism of transcription activation, thereby defining a previously unrecognized mode of transcription activation in cells.

## INTRODUCTION

Bacterial cells are capable of rapidly adapting to different ecological conditions through highly regulated dynamic switching among different gene expression programs, and they do so by selectively activating distinct σ-RNAP holoenzymes(Osterberg et al., 2011). The alternative initiating σ factor, σ^S^ (also known as σ^38^ in *E. coli*) is the master stress regulator that protects many Gram-negative bacteria from detrimental environmental conditions (Lange and Hengge-Aronis, 1991). It also plays indispensable roles in the biofilm formation, virulence, antibiotic tolerance, and antibiotic persistence of human pathogen including *Salmonella enterica*, *Pseudomonas aeruginosa*, as well as pathogenic *E. coli* (Hansen et al., 2008; Murakami et al., 2005; Stewart et al., 2015; Wu et al., 2015). Different stress conditions including antibiotic treatment could strongly induce the transcription and expression of σ^S^ (Battesti et al., 2011), which in turn activates the transcription of ~ 10% genes from the *E. coli* genome by σ^S^-RNAP holoenzyme, thereby rendering bacterial cells resistant to antibiotic treatment and other stresses like hydrogen peroxide, high temperature, low pH, osmotic shock, antibiotic pressure, *etc* (Battesti et al., 2011; Lelong et al., 2007; Weber et al., 2005).

σ^S^ belongs to the group-2 alternative σ factors (Feklistov et al., 2014). The conserved domains of σ^S^ (σ^S^_1.2_, σ^S^_2_, σ^S^_3.1_, σ^S^_3.2_, and σ^S^_4_) interact with RNAP core enzyme through exactly the same interfaces as those of housekeeping σ factor (σ^70^ in *E. coli*) (Liu et al., 2016). σ^S^ shares high degree of protein similarity with σ^70^ and prefers similar sequences at the −10 (TATAAT) and −35 (TTGACA) promoter elements as σ^70^ does(Gaal et al., 2001). Although σ^S^ and σ^70^-regulated genes overlap to some extent, σ^S^ exhibits good selectivity towards its own regulon partially through a “derivation-from-the-optimum” strategy. The σ^S^-RNAP holoenzyme tolerates degenerate promoter sequences (mostly at the −35 element) relatively better than σ^70^-RNAP does, though at the cost of reduced overall transcription activity(Gaal et al., 2001; Maciag et al., 2011; Typas et al., 2007b).

Besides the inferior transcriptional activity of σ^S^-RNAP to σ^70^-RNAP, the cellular amount of *E. coli* σ^S^ is also less than that of σ^70^ in stationary phase and stress conditions (Jishage et al., 1996) and the affinity of σ^S^ is ~ 15 times lower than that of σ^70^ to RNAP core enzyme (Maeda et al., 2000). Therefore, σ^S^ has to cooperate with its allies to compete with σ^70^ for RNAP core enzyme in order to transcribe its own regulon. A large collection of genetic and biochemical data have highlighted the importance of Crl in σ^S^-mediated transcription in bacteria that encode σ^S^ (Cavaliere and Norel, 2016). Crl was demonstrated to directly activate σ^S^-mediate transcription both *in vitro* and *in vivo* (Banta et al., 2013; Banta et al., 2014; Cavaliere et al., 2014; Cavaliere et al., 2015; England et al., 2008; Monteil et al., 2010a; Pratt and Silhavy, 1998; Typas et al., 2007a), and *Crl*-null *Salmonella* and *E. coli* strains displayed impaired biogenesis of curli (important for host cell adhesion and invasion as well as formation of biofilm), increased sensitivity to H_2_O_2_ stress, and reduced virulence due to decrease expression of several σ^S^-regulated genes (Arnqvist et al., 1992; Barnhart and Chapman, 2006; Monteil et al., 2010a; Robbe-Saule et al., 2008; Robbe-Saule et al., 2006).

Crl is a unique transcription activator in bacteria: 1) Unlike other canonical bacterial transcription factors that regulate the activity of the housekeeping σ factor (Browning and Busby, 2016), Crl shows highly stringent specificity to σ^S^ (Banta et al., 2013; Bougdour et al., 2004); 2) Crl broadly activates σ^S^-mediated transcription in a promoter sequence-independent manner(Lelong et al., 2007; Robbe-Saule et al., 2006; Robbe-Saule et al., 2007); and 3) Crl seems to act in at least two stages to boost σ^S^-mediated transcription--the stage of σ^S^-RNAP holoenzyme assembly and the stage of promoter unwinding (Banta et al., 2013; Bougdour et al., 2004; England et al., 2008). Crl has been demonstrated to interact with σ^S^_2_ and probably also RNAP core enzyme(England et al., 2008), but whether or how it interacts with DNA remains elusive. Although crystal and NMR structures of Crl are available(Banta et al., 2014; Cavaliere et al., 2014; Cavaliere et al., 2015), it is still unclear how Crl interacts with σ^S^-RNAP holoenzyme and how such interaction contributes to the transcription activation of σ^S^-RNAP.

RNAP holoenzyme is the bacterial transcription machinery capable of initiating promoter-specific transcription. RNAP holoenzyme binds and subsequently isomerizes double-stranded promoter DNA to form a RNAP-promoter open complex (RPo), a stable intermediate complex competent for initiation of RNA synthesis, through a complicated and multi-step process(Boyaci et al., 2019; Ruff et al., 2015). Conventional bacterial transcription activators such as *E. coli* CAPs (Browning and Busby, 2016; Liu et al., 2017), *C. cresentus* GcrA(Wu et al., 2018), *Chlamydia trachomatis* GrgA(Bao et al., 2012), and Mycobacteria RbpA and CarD (Bae et al., 2015; Hubin et al., 2015) modulate the intermediate steps during RPo transition through simultaneously engaging RNAP holoenzyme and promoter DNA, thereby strengthening the interaction between RNAP and promoter DNA to promote transcription (Browning and Busby, 2016). It is intriguing to know how Crl, as an unconventional transcription activator, activates transcription.

In this study, we have determined a 3.80 Å cryo-EM structure of a transcription activation complex of Crl (*E. coli* Crl-TAC) comprising *E. coli* σ^S^-RNAP holoenzyme, Crl, and a nucleic-acid scaffold mimicking the transcription initiation bubble. In the structure, Crl shields a large solvent-exposed surface of σ^S^_2_, bridges σ^S^_2_ and RNAP-β’ subunit, but makes no contact with promoter DNA. The cryo-EM structure together with data of hydrogen deuterium exchange mass spectrometry (HDX-MS) and mutational studies have converged on a model that Crl activates the σ^S^-RNAP holoenzyme by stabilizing the active conformation of σ^S^ as a chaperon, thereby promoting the interaction between σ^S^ and RNAP or DNA.

## RESULTS

### The cryo-EM structure of *E. coli* Crl-TAC

To understand the structural basis for transcription activation by Crl, we reconstituted the Crl-TAC complex that includes *E. coli* σ^S^-RNAP holoenzyme, Crl, and a nucleic-acid scaffold comprising a 25-bp upstream dsDNA, a 13-bp downstream dsDNA, a non-complimentary transcription bubble (−11 to +2 with respect to the transcription start site at +1), and a 5-nt RNA primer (Fig. 1A and Fig. S1). The cryo-EM structure of *E. coli* Crl-TAC was determined at nominal resolution of 3.80 Å by a single-particle reconstitution method (Table S1 and Fig. S2). The electron density map shows clear signals for the nucleic-acid scaffold, σ^S^, and Crl (Fig. 1B and 1D-E). The crystal structure of Crl could be readily fit into the electron density, suggesting little conformational change of Crl upon interaction with σ^S^-RNAP holoenzyme (Fig. 1D) (Banta et al., 2014).

**Figure 1.**
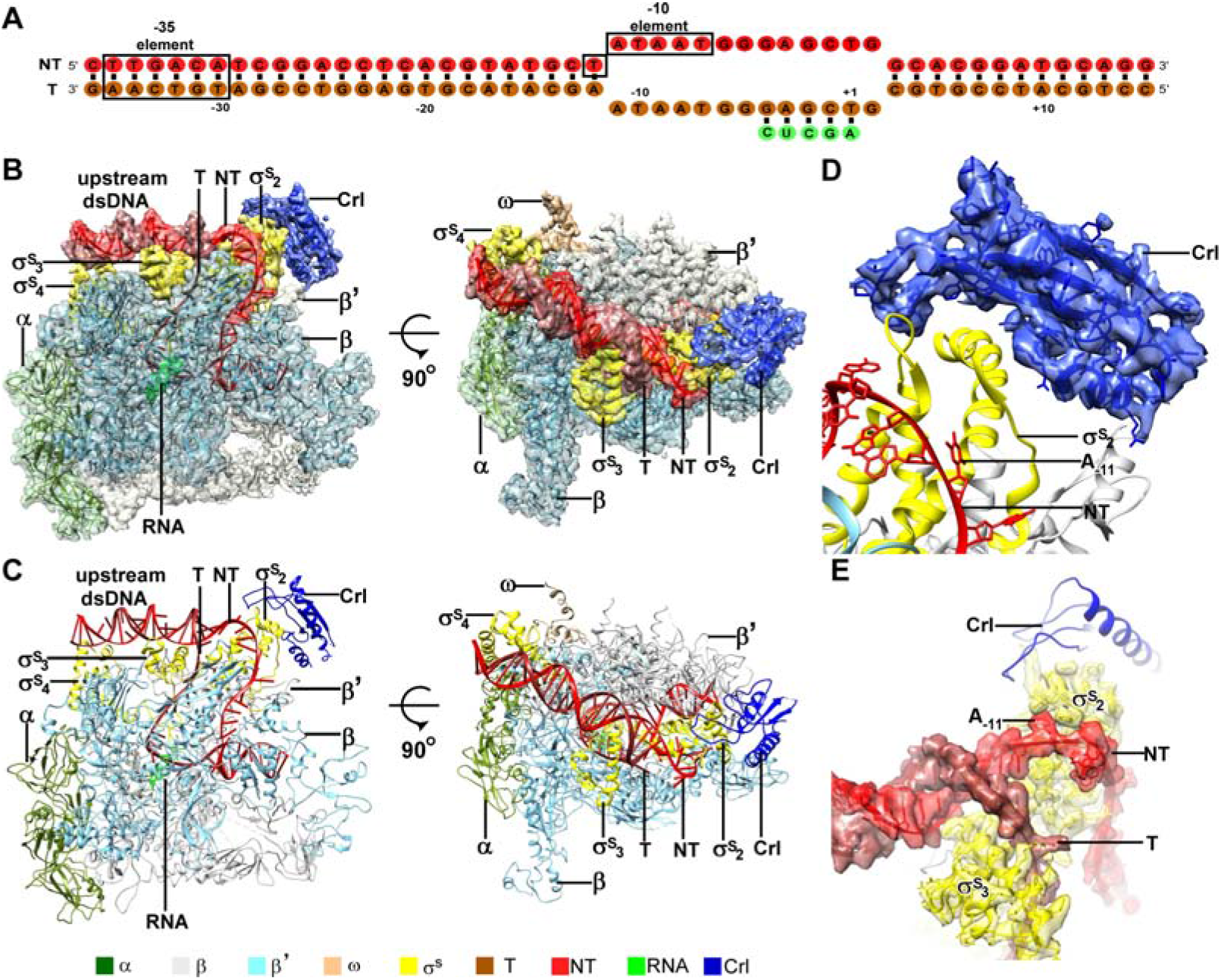
The overall structure of *E. coli* Crl-TAC. **(A)** The scaffold used in structure determination of *E. coli* Crl-TAC. **(B)** The top and front view orientations for cryo-EM electron density map and structure model of *E. coli* Crl-TAC. The RNAP, Crl and nucleic acids are presented as cartoon and colored as in the color scheme. The electron density map is shown as in gray envelop. **(D)** The cryo-EM electron density map (blue transparent surface) for Crl. **(E)** The cryo-EM electron density map (red transparent surface) for the upstream junction of transcription bubble of promoter DNA and σ^S^. NT, nontemplate-strand promoter DNA; T, template-strand promoter DNA.

The cryo-EM structure clearly shows that Crl locates at the outer surface of σ^S^-RNAP holoenzyme (Fig. 1B-C). It mainly interacts with σ^S^_2_ and shields the σ^S^_2_ through an interface of 695 Å^2^; it also contacts a helical domain of RNAP-β’ clamp by a very small interface of ~85 Å^2^ (Fig. 2A-D). Crl approaches to upstream edge of transcription bubble, but makes no direct contact with promoter DNA (Fig. 1D-E and 2A). Such interaction mode between Crl and σ^S^-RPo supports previous findings by genetic, bacterial two-hybrid, cross-linking, SPR, NMR, or bioinformatic techniques (Banta et al., 2013; Banta et al., 2014; Cavaliere et al., 2014; England et al., 2008; Monteil et al., 2010a; Monteil et al., 2010b; Pratt and Silhavy, 1998).

**Figure 2.**
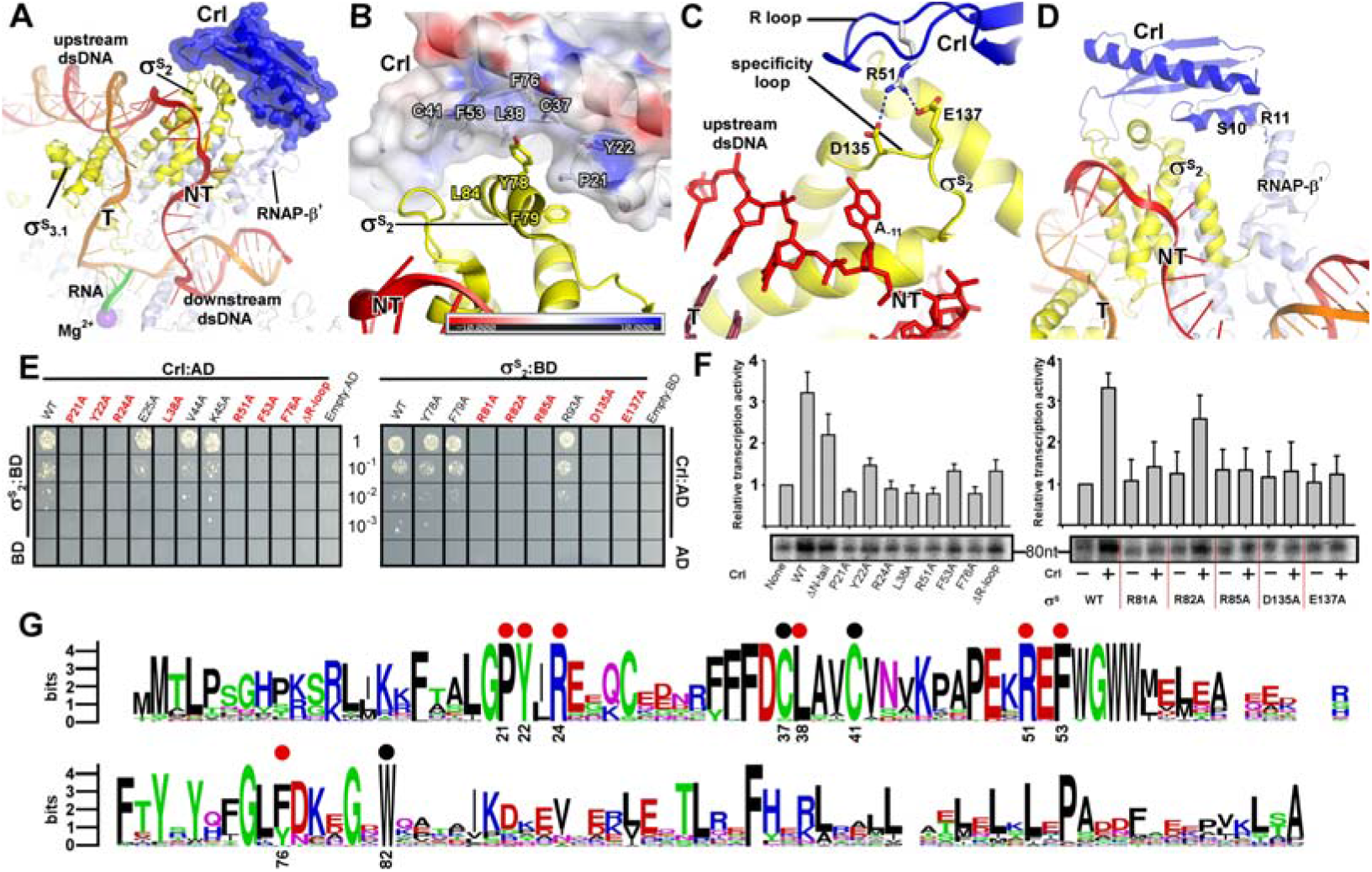
The interactions between Crl and RNAP-holoenzyme. **(A)** Crl binds to the σ^◻^_2_ as well as RNAP-β’ subunit. Crl was represented in blue transparent surface and cartoon. **(B)** σ^S^ is embedded into a shallow hydrophobic groove of Crl. The electrostatic potential surface of Crl was generated using APBS tools in Pymol. **(C)** The detailed salt-bridge interaction between the “R” loopf of Crl and the “specificity loop” of σ^S^. Salt-bridge bonds, blue dash. **(D)** The N-terminal tail of Crl makes potential weak interactions probably through R11 and S10 with RNAP-β’ subunit. The colors are as in Figure 1. **(E)** The yeast two-hybrid assay reveals key interface residues (red) of Crl (left) or σ^S^ (right). Mutating the key residues disrupts interactions between Crl and σ^S^. **(F)** The *in vitro* transcription assay shows that mutating most of key interface residues of Crl (left) substantially impairs its transcription activation activity, and mutating most key interface residues of σ^S^ resulted in reduced response to Crl. ΔR-loop, replacing residue 43-51 with a “GSGS” linker. **(G)** Protein sequence alignment of Crl from 72 non-redundant bacterial species. Filled circles indicate residues involved with interactions with σ^S^; filled red circles indicate key contact residues. The residues are numbers as in *E. coli* Crl. NT, nontemplate-strand promoter DNA; T, template-strand promoter DNA.

### Crl interacts with σ^S^ and RNAP core enzyme

In the structure of *E. coli* Crl-TAC, helix α2 of σ^S^_1.2_ (residues 72-82) and σ^S^_NCR_ (residues 83-89) are embedded into a shallow groove on Crl and the nearby “specificity loop” of σ^S^_2.3_ (residues 133-141) is further anchored by Crl (Fig. 2A-C, S3 and S4). Several Crl residues (P21, Y22, C37, L38, C41, F53, F76, and W82) create a hydrophobic patch in the shallow groove and make Van del Waals interactions with residues (Y78, F79, R82, L84, and R85) of σ^S^ (Fig. 2B and Fig. S3B). Moreover, potential polar interactions between Crl (R24) and σ^S^ (D87) might also contribute to the interaction. The interface residues identified here recapitulate most hits in a previous genetic screening for identification of interacting residues between σ^S^ and Crl (Banta et al., 2014). Intriguingly, most evolutionarily conserved residues of Crl are clustered in the shallow groove, implicating a functional relevance of its interaction with σ^S^ (Fig. 2G). Moreover, the Crl derivatives F53A and W82A posed salmonella cells more sensitive to H_2_O_2_ stress (Monteil et al., 2010a), demonstrating the physiological importance of such interface.

We subsequently evaluated contribution of each interface residue to σ^S^-Crl interaction using a yeast two-hybrid assay. The results show that mutating most interface residues (P21W, Y22A, R24A, L38A, F53A, or F76A of Crl; R81A, R82A or R85A of σ^S^_2_) substantially impaired σ^S^-Crl interactions (Fig. 2E). Moreover, the results from *in vitro* transcription assay show that alanine substitutions of R81, R82, or R85 of σ^S^ impeded its response to Crl; and alanine substitutions of L38 or F53 of Crl also impaired its ability of transcription activation (Fig. 2F). The results validate our structure and justify the significance of the σ^S^-Crl interface for the transcription activation activity of Crl.

Previous study suggested that a conserved “DPE” motif on σ^S^_2_ plays an indispensable role in its interaction with Crl (Banta et al., 2013; Banta et al., 2014). The “DPE” motif is located on the “specificity loop” of σ^S^_2_, an essential structural element responsible for recognizing and stabilizing the unwound nucleotide at the most conserved position of promoter DNA (*i.e.* position −11 of σ^S^ and σ^70^) in all bacterial transcription initiation complexes (Campagne et al., 2014; Li et al., 2019; Lin et al., 2019; Liu et al., 2016; Zhang et al., 2012). In our structure, the “specificity loop” encloses the NT-11A nucleotide as reported (Fig. 2C and 4D) (Liu et al., 2016). Notably, the conformation of “specificity loop” is secured by the “R” loop (residues 41-53) of Crl. The side chain of Crl R51 reaches D135 and D137 of the conserved “DPE” motif of σ^S^_2_, and probably makes salt-bridge interactions with the two aspartic acids (Fig. 2C and S3C). Our results of yeast two-hybrid and *in vitro* transcription assays show that mutating either D135/D137 of Crl or R51 of σ^S^_2_ significantly compromised the Crl-σ^S^_2_ interaction and Crl-mediated transcription activation, highlighting the importance of such interface (Fig. 2E-F).

Our structure of Crl-TAC also explains the strict specificity of Crl to σ^S^. Crl binds to two of the least conserved regions of bacterial σ factors--the σ^S^_NCR_ and the “specificity loop” (Fig. S4A). The sequence alignments of 10 bacterial σ^S^ and 10 primary σ factors clearly show that most Crl-interacting residues on σ^S^ are not present in σ^70^ (Fig. S4B-C).

In agreement of the previous SPR results (England et al., 2008), the electron density of Crl also suggest possible interactions between Crl and the RNAP-β’ clamp domain (Fig. 1D). The interface is relatively small and only involves residues S10 and R11 of the Crl N-terminal loop (Fig. 2D). Deletion of the N-terminal loop of Crl only shows marginal effect on its activity of transcription activation, suggesting a dispensable role of such interface (left panel of Fig. 2F).

### Crl facilitates assembly of σ^S^-RNAP holoenzyme by stabilizing σ^S^

It has been suggested that Crl may function as a σ^S^ chaperon to facilitate the assembly of σ^S^-RNAP holoenzyme (Banta et al., 2013). We confirm that Crl is able to increase ~ 3.8 fold of the binding affinity between σ^S^ and RNAP core enzyme in a fluorescence polarization assay using fluorescein-labeled σ^S^ (Fig. 3A). Moreover, an “R loop”-mutated derivative of Crl completely lost its effect on the assembly of σ^S^-RNAP holoenzyme, while deleting the N-terminal tail of Crl barely affects the assembly (Fig. 3B), in agreement with our *in vitro* transcription results (Fig. 2E-F). Such results further suggest that Crl facilitates the assembly of σ^S^-RNAP holoenzyme mainly through its interaction with σ^S^ rather than through its interaction with RNA-β’ clamp.

**Figure 3.**
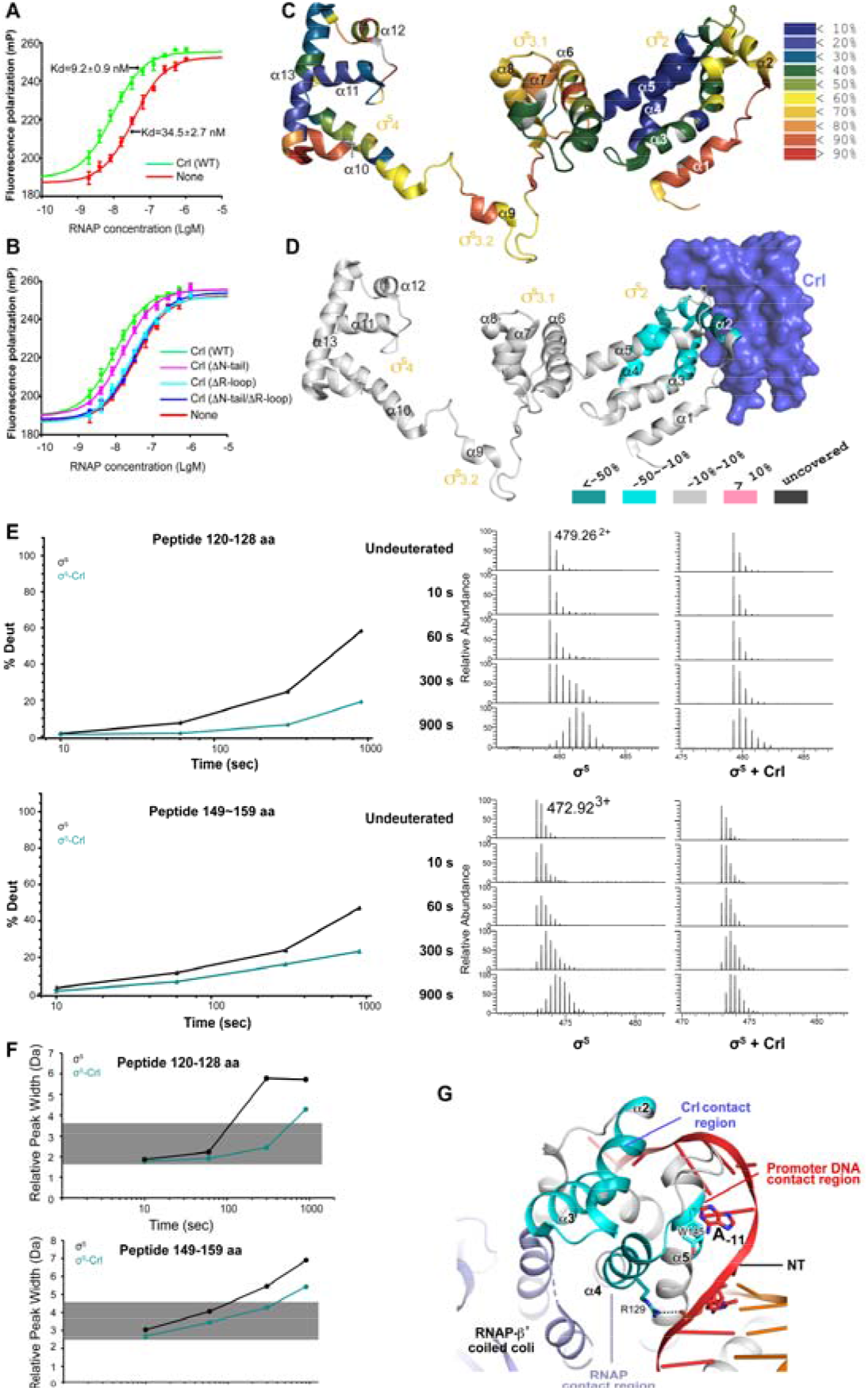
Crl promotes assembly of σ^S^-RNAP holoenzyme by stabilizing σ^S^_2_. **(A)** Crl increases binding affinity between RNAP and σ^S^ in a FP assay. **(B)** Deletion the “R” loop but not N-tail (ΔN-tail; deleting residue 1-11) abolishes the ability of Crl on assembly of σ^S^-RNAP holoenzyme. (**C**) HDX Profile of free σ^S^ after 10 s deuteron. Incorporation was mapped onto the cryo-EM structure of σ^S^. The inset shows the color coding for different percentages of deuteron incorporation. (**D**) HDX changes for σ^S^ in the presence of Crl as compared to free σ^S^, mapped on the cryo-EM structure of σ^S^. The percentage difference in deuterium uptake values represents an average across all four time points ranging from 10 s to 900 s. For heatmap color coding, pink, cyan, and gray indicates increase, decrease, and no significant changes of HDX, respectively. Dark grey represents regions that were not consistently resolved in all HDX experiment. (**E**) Deuterium uptake plots and mass spectra of indicated peptides from helices α4 and α5 in the absence and presence of Crl. Left: The deuterium uptake data are plotted as percent deuterium uptake versus time on a logarithmic scale. Right: Mass spectra of indicated peptides at different labeling time points, with the mass spectra of undeuterated samples shown as controls. (**F)** Peak-width analysis at different time points for peptides from helices α4 and α5 reveal the existence of EX1 kinetics in the absence of Crl. (**C-F**) The HDX experiments were repeated at least twice. (**G**) A highlight of σ^S^_2_ regions with reduced HDX rate upon Crl binding, the colors are as in (D).

**Figure 4.**
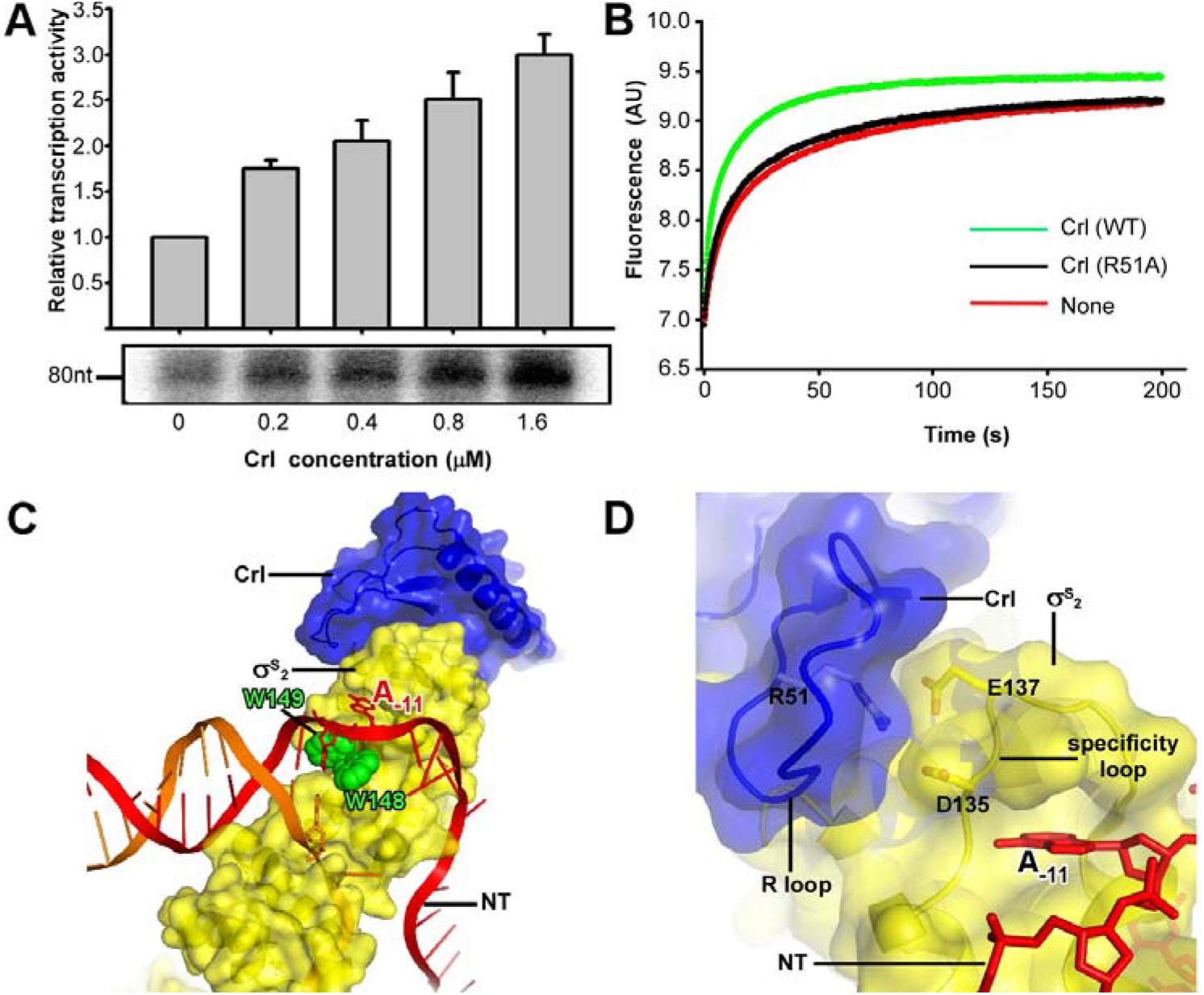
Crl facilitates promoter unwinding. **(A)** Crl increases transcription by a pre-assembled σ^S^-RNAP holoenzyme from P*osmY* promoter in a concentration dependent manner. **(B)** Crl increases the rate of equilibration of σ^S^-RPo formation in a fluorescence stopped-flow experiment while mutating R51 of Crl “R” loop diminished the effect of Crl on RPo formation. **(C)** The upstream promoter DNA is unwound by the W-dyad (W148 and W149) and the unwound A_−11_ nucleotide of the nontemplate strand DNA is recognized and stabilized by a protein pocket on σ^S^_2_. **(D)** The “R” loop of Crl stabilizes conformation of “specificity” loop that forms the pocket for unwinding and recognizing the A_−11_ nucleotide of nontemplate strand promoter DNA.

Previous report also showed that Crl shifts the equilibrium towards σ^S^-RNAP holoenzyme formation rather by promoting the association between σ^S^ and RNAP than by preventing the dissociation of the two (England et al., 2008). Such facts imply that Crl may stabilize σ^S^ in a conformation that is more accessible for RNAP core enzyme. To investigate into this hypothesis, we characterized the in solution dynamic conformations of full-length σ^S^ using HDX-MS.

Deuterium exchange of σ^S^ was first monitored in the absence of Crl at different time points. At the earliest time point 10 s, low deuterium incorporation (≤50 %) was observed for most peptides from σ^S^_2_ and σ^S^_4_(Fig. 3C and Fig. S5), and the deuterium uptake of these peptides increases gradually over time (Fig. S5 and Fig. S6–S7). Such facts indicate that these two domains generally adopt folded structures in solution, as observed in the cryo-EM structure. Meanwhile, although σ_3.1_ appears to be folded in our cryo-EM structure (helix α6-α8), most peptides from these regions, except for those ranging residues 182-197, manifest high deuterium incorporation level (≥50 %) at the earliest time point 10 s and unchanged deuterium uptake up to 900 s (Fig. 3C and Fig. S5, S7), indicating that σ_3.1_ is largely solvent-exposed in solution. Although domain σ_1.1_ (residues 1-55) was not resolved in the cryo-EM structure, good peptide coverage was achieved for this region in HDX-MS experiment and shows rapid deuterium incorporation (Fig. S8). Consistent with their remote locations in the cryo-EM structure, helix α1 from σ_1.2_ (residues 56-67) and σ_3.2_ linker (residues 218-245) generally appear to be solvent-exposed in HDX-MS experiment (Fig.3C and Fig.S5, S7). The HDX profile of σ^S^ in the absence of Crl also agrees well with a recent report (Cavaliere et al., 2018).

We then monitored the deuterium exchange profile of σ^S^ in the presence of Crl. From HDX-MS results, increased protection from HDX was consistently observed for peptides ranging residues 74-99 (Fig. 3D, and Fig. S6A-B), which span residues 74-82 from σ^S^_1.2_, σ^S^_NCR_ (residues 83-89) and its flanking sequence. Collectively, the decreased deuteron uptakes in this region of σ^S^_2_ agree very well with the cryo-EM structure (Fig. 3D, and Figure 2A-B). Moreover, increased protection from HDX was also observed for peptides from α4 (ranging 122-134aa) and α5 (ranging residues 145-148aa), all of which locate far away from the interface between σ^S^ and Crl as observed in the cryo-EM structure (Fig. 3D). Such fact suggests an allosteric stabilizing effect of Crl on σ^S^, and we thus analyzed the raw spectra of all peptides from these two regions.

To our surprise, in the absence of Crl, typical bimodal shaped isotope clusters were observed for all peptides spanning 122-131aa and 148-157aa at longer time points (Fig. 3E, and Fig. S6C-D). Consistently, maximum peak widths of these peptides also increased dramatically as the labeling prolongs (Fig.3F), indicating an EX1 exchange mechanism in these regions(Weis et al., 2006). Together, these evidences suggest that, in the absence of Crl, part of these two regions underwent an unfolding event that is long enough for the hydrogen-to-deuterium exchange to happen at all exposed residues(Wales and Engen, 2006). Addition of Crl apparently helps to restrain the two regions on σ^S^_2_ in their low HDX conformation (Fig.3E-F and Fig. S6C-D), and the HDX protection effect seems to be highly pronounced for peptides from α4.

Notably, α4 is one of the major anchor point on σ^S^_2_ for RNAP-β’ subunit (Fig. 3G). Hence, the stabilizing effect rendered on helix α4 by Crl probably facilitates the docking of σ^S^_2_ to RNAP-β’ subunit for subsequent proper positioning of other domains into the RNAP core enzyme, thereby promoting the assembly of functional σ^S^-RNAP holoenzyme. Such hypothesis is also supported by the increased rate of association during formation of σ^S^-RNAP holoenzyme in the presence of Crl (England et al., 2008).

### Crl stabilizes σ^S^_2_ to assist in promoter unwinding

Previous report suggested that Crl is also able to boost σ^S^-mediated transcription at a step after σ^S^-RNAP holoenzyme assembly (Bougdour et al., 2004; England et al., 2008). We here show that Crl increases transcription activity of a pre-assembled σ^S^-RNAP holoenzyme in an *in vitro* transcription assay (Fig. 4A), further supporting the previous finding.

To understand how Crl activates σ^S^-mediated transcription beyond assembly of σ^S^-RNAP holoenzyme, we modified a stopped-flow fluorescence assay to monitor the potential effect of Crl on promoter unwinding by σ^S^-RNAP holoenzyme. Such technique has been employed to study the kinetics of promoter unwinding by *E. coli* σ^70^-RNAP holoenzyme (Sullivan et al., 1997). The σ^S^-RNAP holoenzyme alone readily unwinds the prompter DNA and slowly reaches equilibration of RPo formation after 100 seconds; importantly, Crl increase the rate of equilibration of σ^S^-RPo formation (faster to reach the plateau; Fig. 4B), demonstrating that Crl also functions at the step of promoter unwinding.

A key structural element in σ^S^, i.e. the “specificity loop”, interacts the first unwound nucleotide and thus stabilizes transcription bubble during the following steps of transcription initiation (Fig. 4C-D). In our structure, Crl uses its residue R51 to interact with D135 and D137 from the “specificity loop” and secures the latter in a conformation that readily accepts the unwound nucleotide. We thus propose that Crl may contribute in the promoter unwinding step by stabilizing the “specificity loop” in its active conformation. Notably, alanine substitution of Crl residue R51 abolishes the effect of Crl on promoter unwinding (Fig. 4B), further supporting such hypothesis.

Meanwhile, residues from α4 and α5 that showed decreased HDX rate in the presence of Crl also form intimate interaction with the upstream junction of promoter DNA (Fig. 3G). In the cryo-EM structure, residue R129 from α4 may salt-bridge with the DNA phosphate backbone (Fig.3G). Meanwhile, residue Y145 from α5 sits just underneath the base of the unwound A_−11_ (Fig.3G), and might facilitate the flipping of A_−11_ by pi-stacking with its base. Such facts thus suggest that the rigidifying effect rendered on helices α4 and α5 by Crl might also facilitate the docking of promoter DNA and indirectly contribute to the unwinding process.

Together, our biochemical data, cryo-EM structure and HDX-MS results clearly support a model where Crl functions as a stabilizing chaperon to facilitate the assembly of σ^S^-RNAP holoenzyme and the promoter unwinding.

## DISCUSSION

Crl has been discovered as the specific transcription activator of σ^S^-RNAP holoenzyme about 20 years ago. A large collection of biochemical, biophysical and genetic data implies that Crl may activate transcription in an unprecedented manner. In this study, we determined a 3.80 Å cryo-EM structure of *E. coli* Crl-TAC (Fig. 1). The structure shows that Crl shields a large and otherwise solvent-exposed surface on σ^S^_2_ in σ^S^-RNAP holoenzyme, stabilizes the key “specificity loop” of σ^S^_2_, but doesn’t contact promoter DNA (Fig. 2). Subsequent HDX-MS results recapitulate the interaction between Crl and the α2 helix of σ^S^_2_ in solution and further unravel that the stabilizing effect of Crl on σ^S^_2_ extends beyond the helix α2 to α4 and α5 (Fig. 3C-F). Considering the role of α4 and α5 of σ^S^_2_ in anchoring RNAP-β’ subunit and promoter DNA, our cryo-EM structure and HDX-MS data thus point to a model where binding of Crl stabilizes the conformation of σ^S^ to promote the assembly of σ^S^-RNAP holoenzyme and the unwinding of promoter DNA.

Most bacterial transcription factors are capable of making interactions simultaneously with promoter DNA and RNAP and create a direct linkage between them; while the transcription activators in eukaryotic systems typically interact with both enhancer DNA element and mediator proteins, the latter of which bridge various transcription activators to transcription core machinery(Jeronimo and Robert, 2017). Thereby, it is widely accepted that transcription factors activate transcription by enhancing the interaction between promoter DNA and the transcription machinery (Browning and Busby, 2016). The most intriguing question is how Crl activates transcription without interaction with promoter DNA.

Our structure reveals that Crl activates transcription through stabilizing the critical structural elements of σ^S^_2_—the “RNAP-anchoring helix” and “specificity loop”. The effect on the “specificity loop” is straightforward as Crl makes direct interactions with it. It is intriguing that the conformations of “RNAP-anchoring helix” is remotely restrained by Crl binding. Although it is unknown how exactly the signal of Crl binding is allosterically transmitted to the “RNAP-anchoring helix”, we infer that Crl probably clamps the two helices of σ^S^_1.2_ and σ^S^_NCR_, reduces the internal motion of the helix bundle, and consequently stabilizes the conformer competent for engagement to RNAP.

In contrast to canonical bacterial transcription activators that interact with the RNAP-α subunit and/or region 4 of σ factor, Crl specifically activates the σ^S^-regulated genes by making interactions with region 2 of σ^S^. Such mode of interaction has also been employed by a few recently discovered transcription activators, such as *C. crescentus* GcrA, *Mycobacterium tuberculosis* RbpA, and *Chlamydia trachomatis* GrgA, suggesting that the region 2 of σ factor could also serve as a hub for docking various transcription activators.

Collectively, here we revealed that Crl increases transcriptional activity of σ^S^-RNAP in a DNA contact-independent manner and through stabilizing the key structural elements of σ^S^. Our study provides the structural basis and molecular mechanism of an unprecedented example of transcription activation by *E. coli* Crl. The combined effect of Crl--facilitating assembly of σ^S^-RNAP holoenzyme and assisting promoter unwinding--would help σ^S^ to outcompete the housekeeping σ and substantially increase the transcription activity of σ^S^-RNAP holoenzyme for expression of stress-related genes. The unique DNA contact-independent mechanism also provides a new paradigm for bacterial transcription activation.

## STAR*METHODS

Detailed methods are provided in the online version of this paper and include the following:

KEY RESOURCES TABLE
CONTACT FOR REAGENT AND RESOURCE SHARING
EXPERIMENTAL MODEL AND SUBJECT DETAILS
METHOD DETAILS

Plasmid construction
Protein preparation
Nucleic-acid scaffolds.
**Complex reconstitution of *E. coli* Crl-TAC**
**Cryo-EM structure determination of *E. coli* Crl-TAC**
**Hydrogen–deuterium exchange mass spectrometry (HDX-MS) of σ^S^**
Yeast two-hybrid assay.
Fluorescence labeling
Fluorescence polarization (FP) assay
Stopped-flow assay
*In vitro* transcription assay
**Quantification and statistical analysis**
DATA AND SOFTWARE AVAILABILITY

## SUPPLEMENTAL INFORMATION

Supplemental Information including 5 figures and 2 tables can be found with this article online at XXXX.

## ACKNOWLEDGEMENTS

The Work was supported by the National Key Research and Development Program of China (2018YFA0900700), the Strategic Priority Research Program of the Chinese Academy of Sciences (XDB29020000), the National Natural Science Foundation of China (31822001), and the Leading Science Key Research Program of CAS (QYZDB-SSW-SMC005). We thank Dr. Zhaocai Zhou for generous gift of pET28a-TEV plasmid, Dr. Shenghai Chang at the cryo-EM center of Zhejiang University for assistance of grid preparation and data collection. We thank the state key laboratory of bioorganic and natural products chemistry at Shanghai institute of organic chemistry at CAS for sharing the stopped-flow fluorescence spectrometer.

## AUTHOR CONTRIBUTIONS

J. X. performed biochemical experiments. K. C. performed the HDX experiments. J. X. and L. S. solved the structure. J. S., L. Y. and C. F. assisted in cryo-EM data collection and structure determination. L. L. assisted in stopped-flow assay. G. Z, B. Y., Y. F., and Y. Z. designed experiments, analyzed data, and wrote the manuscript.

## SUPPLEMENTAL FIGURES

**Fig S1.**
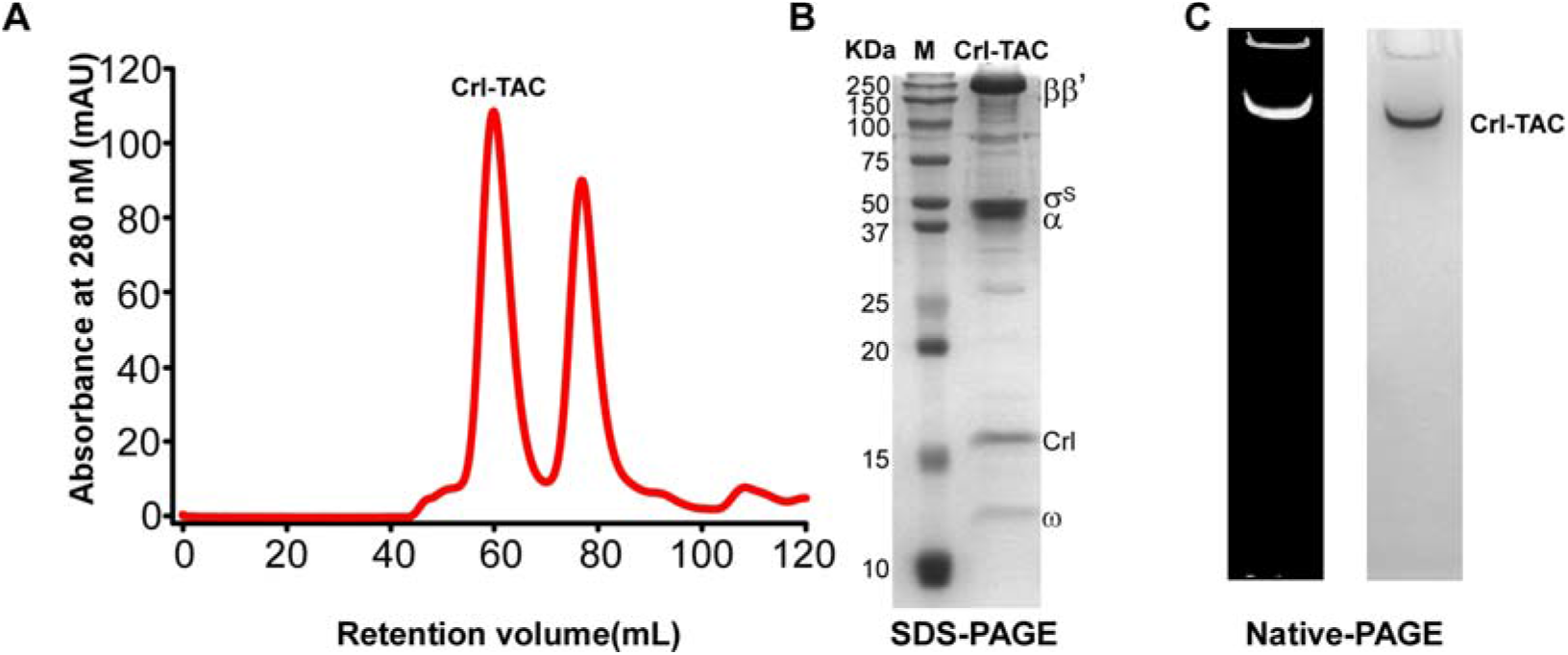
The preparation of *E. coli* Crl-TAC complex. **(A)** Elution peaks of *E. coli* Crl-TAC from a size-exclusion column. **(B)** The SDS-PAGE and **(C)** the native-PAGE of the *E. coli* Crl-TAC. The gel was first stained with SYBR Gold for nucleic acids and then with Coomassie Brilliant Blue for proteins.

**Fig. S2.**
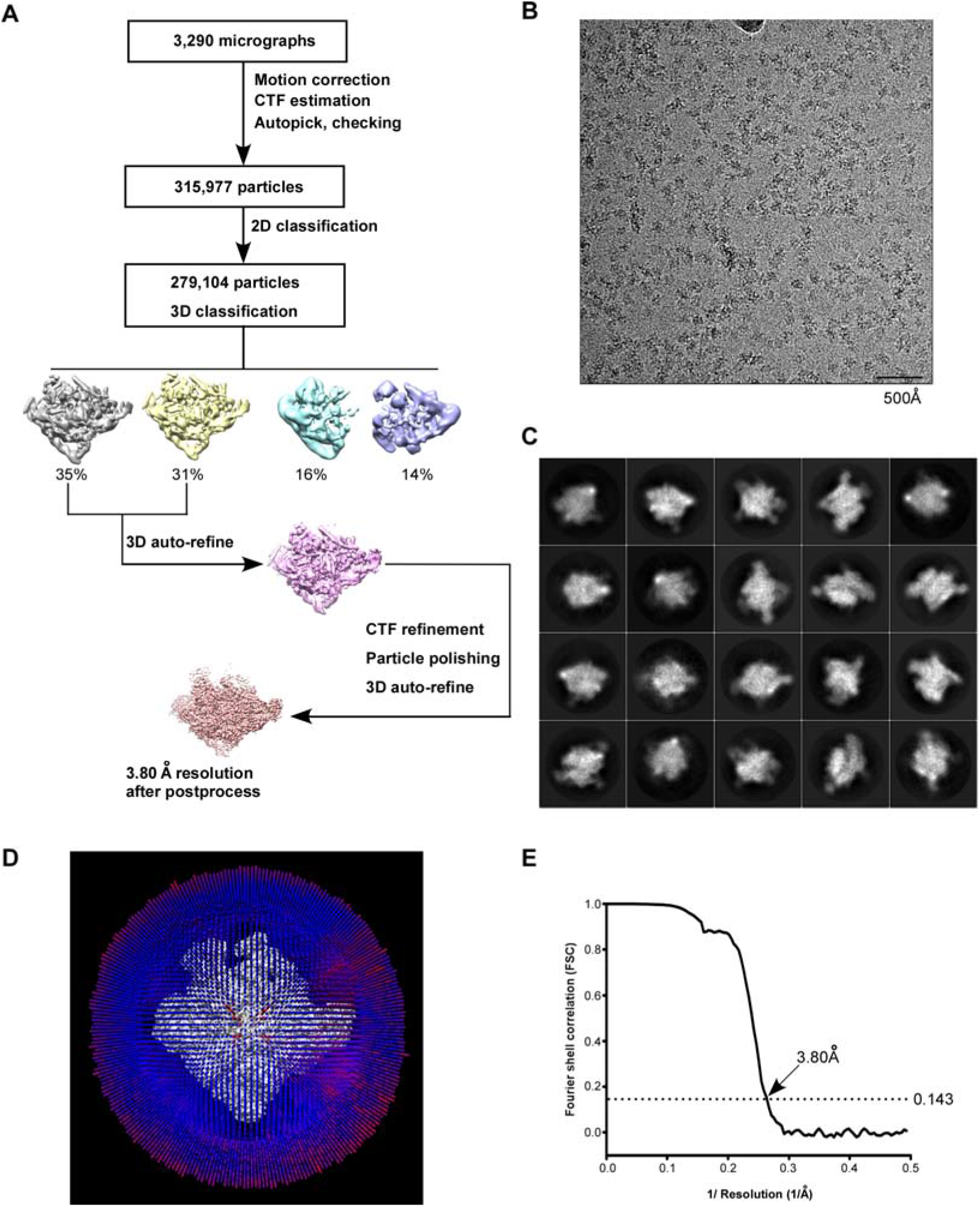
The processing pipeline for cryo-EM map construction of *E. coli* Crl-TAC. **(A)** The flowchart of data processing for cryo-EM structure of *E. coli* Crl-TAC. **(B)** The representative cryo-EM micrograph. **(C)** The representative 2D classifications of *E. coli* Crl-TAC single particles. **(D)** The angular distribution of *E. coli* Crl-TAC particle projections. (E) The gold-standard FSC of *E. coli* Crl-TA. The dotted line represents the 0.143 FSC cutoff, which indicates a nominal resolution of 3.80 Å.

**Fig. S3.**
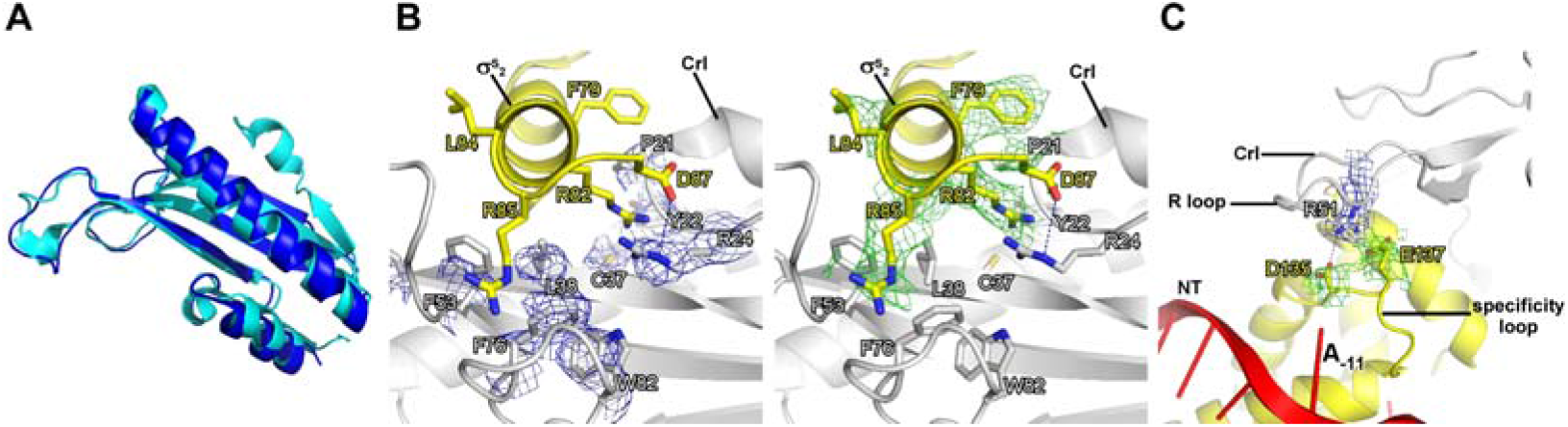
The detailed interactions between Crl and σ^S^_2_. **(A)** Structure superimposition of Crl proteins from crystal structure of Crl (PDB: 4Q11; cyan) and Crl in *E. coli* Crl-TAC (blue). **(B)** The σ^S^_2_ inserts into a hydrophobic groove on surface of Crl. **(C)** The interaction between the “R” loop of Crl and the “specificity” loop of σ^S^_2_. The cryo-EM electron density maps for the side chains of interface residues of Crl and σ^S^_2_ are shown as in blue and green mesh, respectively. The C, N, and O atoms of Crl residues are shown in white, blue, and red, respectively. The C, N, and O atoms of σ^S^ residues are shown in yellow, blue, and red, respectively.

**Fig. S4.**
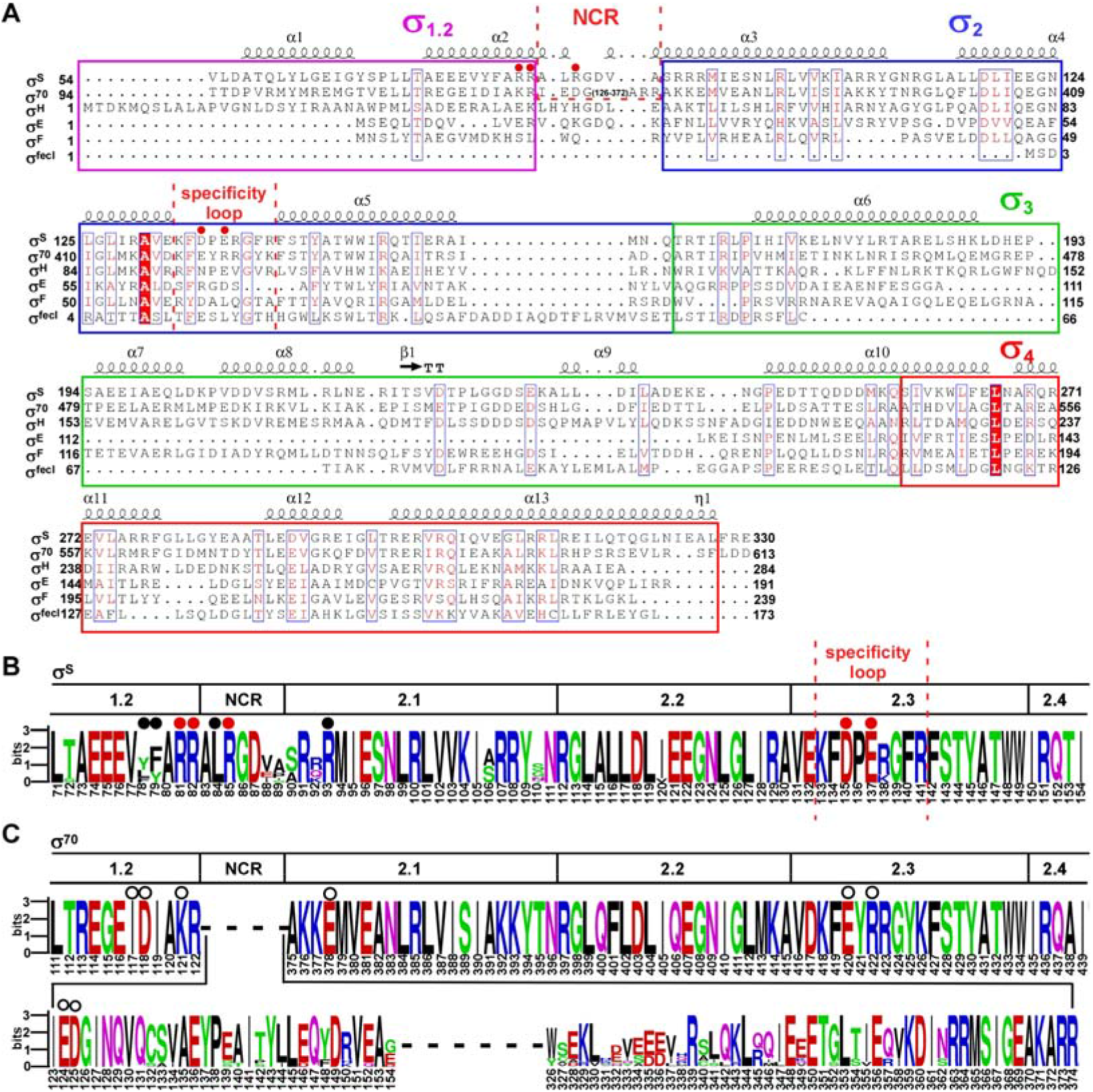
The sequence alignment of bacterial sigma factors. **(A)** Sequence alignment of six σ^70^-type *E. coli* σ factors. **(B)** The sequence alignment of σ^70^ of ten representative bacterial species (*Proteus mirabilis, Escherichia coli, Salmonella enterica* serovar Typhi, *Yersinia enterocolitica, Shigella flexneri, Vibrio cholerae, Dickeya dadantii, Providencia stuartii, Xenorhabdus bovienii, Photobacterium profundum*). **(C)** The sequence alignment of σ^S^ of the above ten bacterial species. The residues are numbered as in *E. coli*.

**Fig. S5.**
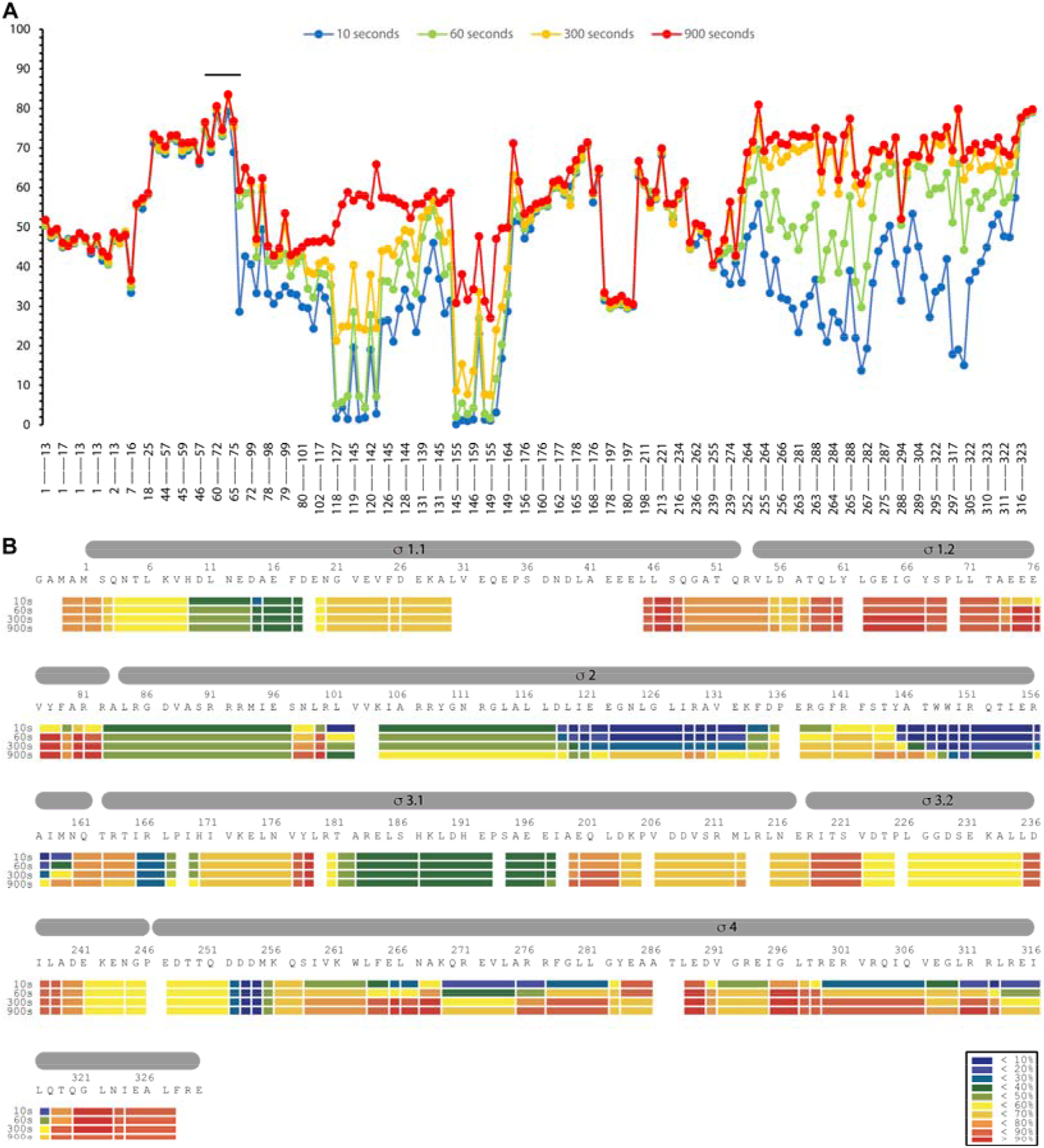
HDX profile of the σ^S^ protein in the absence of Crl. **(A)** The relative deuteration levels (%D) of each peptide were calculated with the assumption that a fully deuterated sample retains 90% D in current LC setting and plotted as a function of peptide position. The blue to red lines represents data acquired at 10 sec up to 900 sec deuteration. (**B**) heat map of **σ**^S^ protein at different time points. The inset shows the color coding for different percentages of deuteron incorporation.

**Fig. S6.**
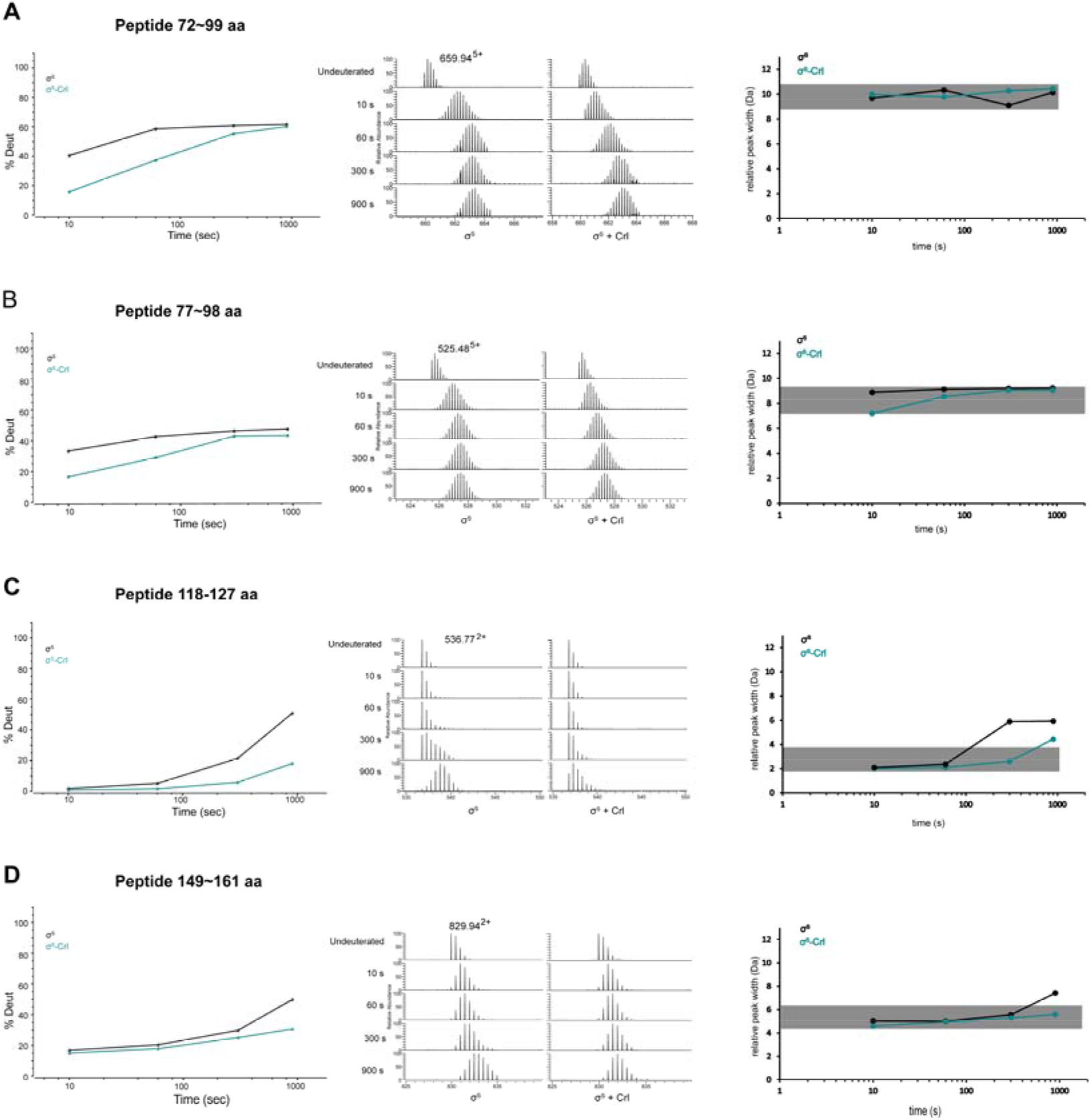
Representative peptides from helix α2, α3, α4 and α5 of σ^S^_2_ whose hydrogen-to-deuterium exchange rate decreased considerably in the presence of Crl. (a-c) Deuterium uptake plots, peak width plots and mass spectra of indicated peptides from σ^S^_2_ in the absence and presence of Crl. Left: the deuterium uptake data are plotted as percent deuterium uptake versus time on a logarithmic scale; middle: mass spectra of indicated peptides at different labeling time points, with the mass spectra of undeuterated samples shown as controls; right: peak-width analysis at different time points for peptides covering helices α2-α5.

**Fig. S7.**
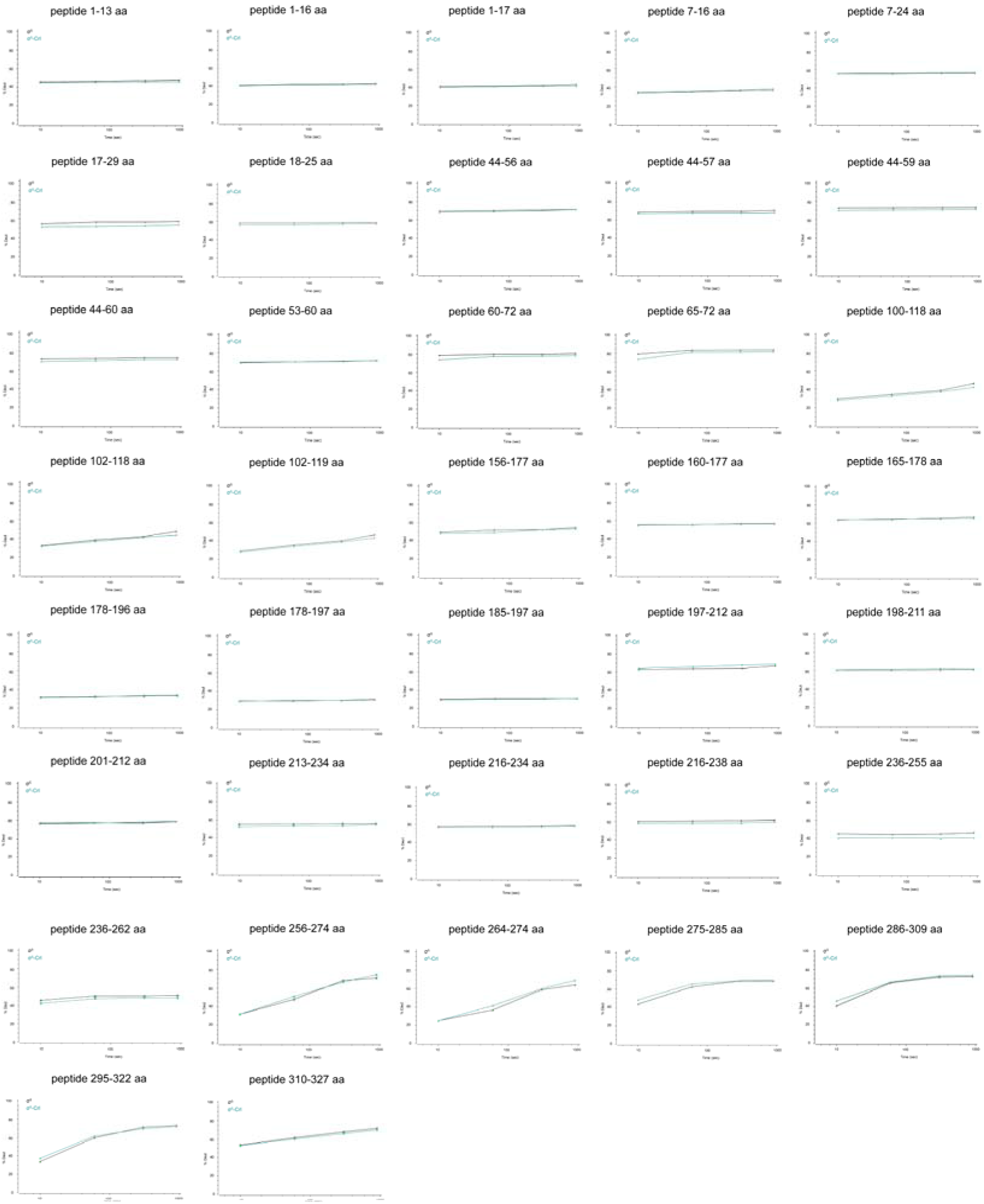
Deuterium uptake plots of additional peptides from σ^S^ whose hydrogen-to-deuterium exchange rate did not change in the presence of Crl. The deuterium uptake data are plotted as percent deuterium uptake versus time on a logarithmic scale.

**Fig. S8.**
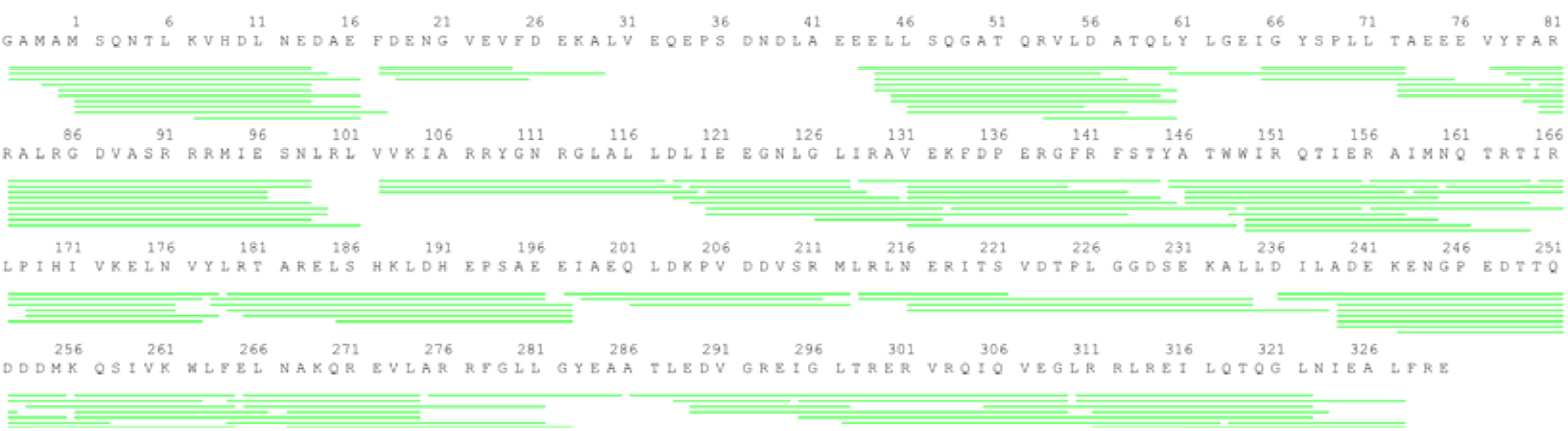
Peptide coverage of σ^S^. The sequence coverage map for **σ**^s^ in HDX-MS study. Green lines shown below the protein sequence represent the digested peptides identified and analyzed in this study.

## STAR*METHODS

**Table S1.**
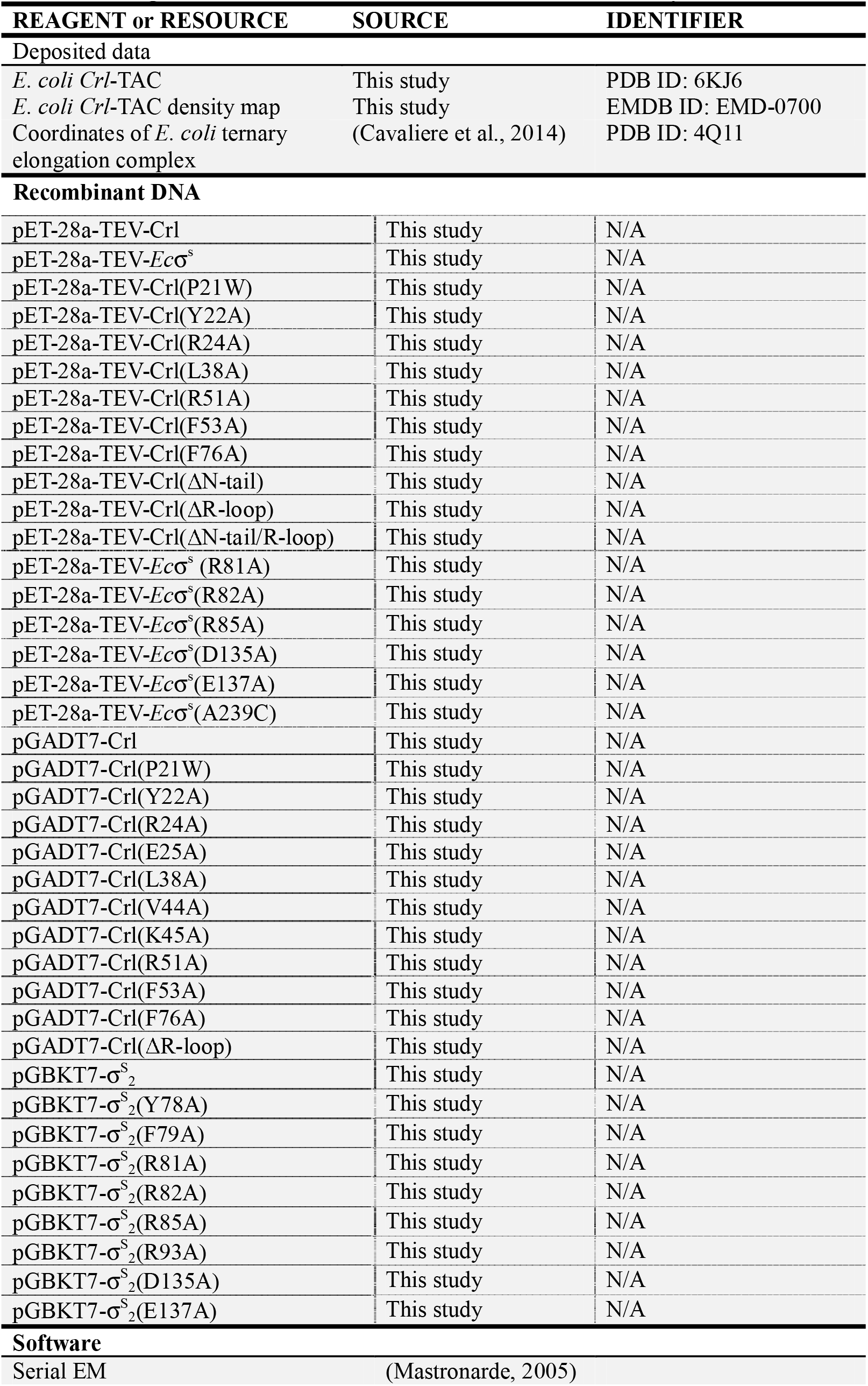
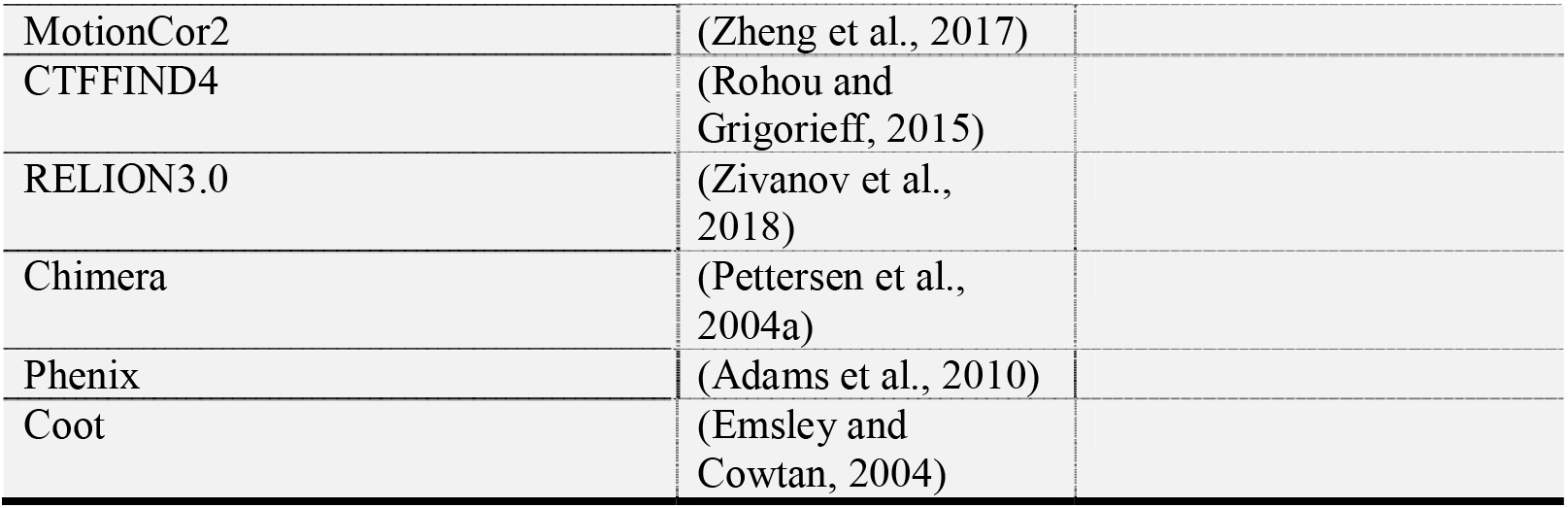
Reagents, resources, DNA, and software used in this study.

**Table S2.**
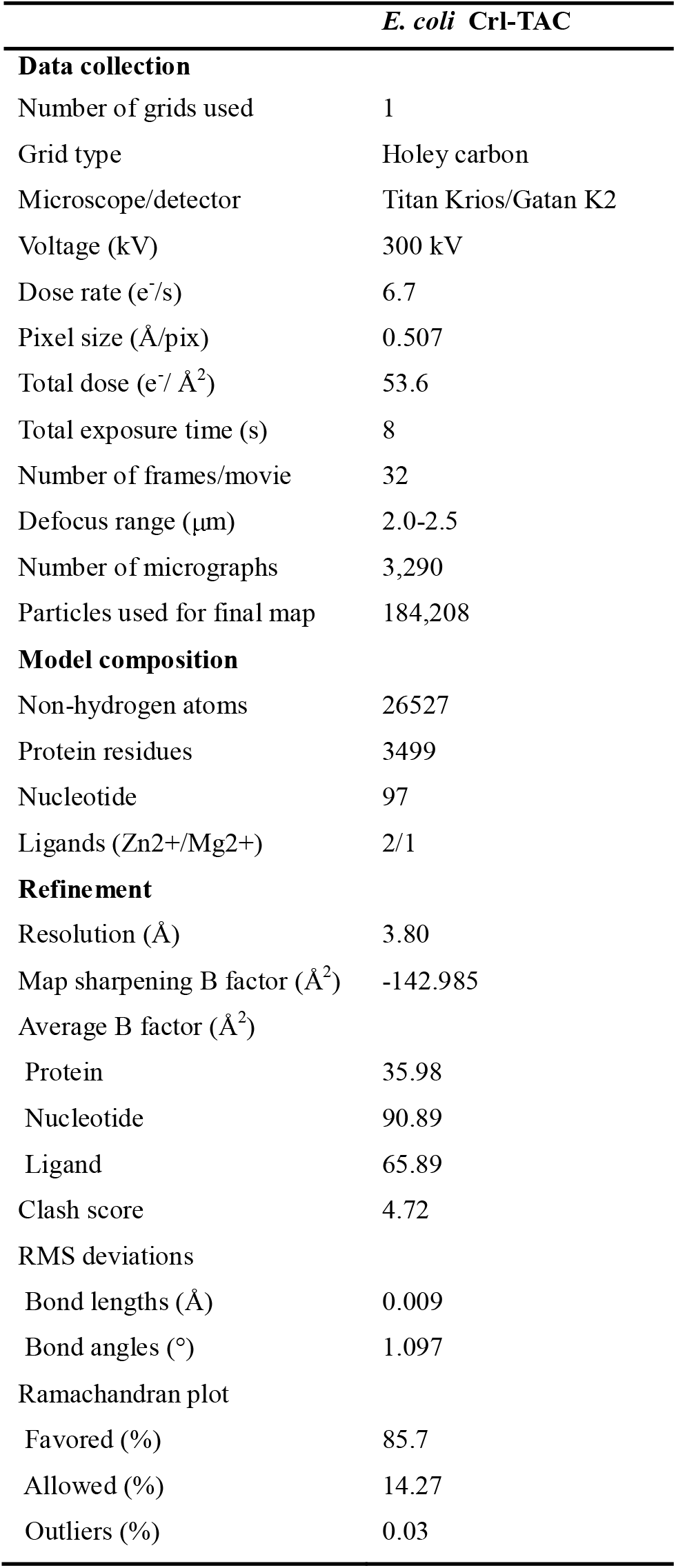
The statistics of the cryo-EM structure of *E. coli* Crl-TAC.

## METHOD DETAILS

### Plasmid construction

DNA fragments containing *E. coli crl* and *rpoS* were amplified from *E. coli* genomic DNA and cloned into pET28a-TEV using the Ezmax one-step cloning kit (Tolo Bio-tech, China.). Point mutations of *crl* or *rpoS* were generated through site-directed mutagenesis (Transgen biotech, Inc.). pET-28a-TEV-Crl(ΔR-loop) was constructed by replace residues 43-51 of Crl by a ‘GSGS’ linker.

pGADT7-Crl and pGBKT7-σ^S^_2_ were constructed using the Gateway LR clonase II (Invitrogen, Inc.). The derivatives of pGADT7-Crl and pGBKT7-σ^S^_2_ were generated through site-directed mutagenesis.

### Protein preparation

The *Ec* RNAP core enzyme was over-expressed and purified from *E. coli* BL21(DE3) carrying p*Ec*ABC and pCDF-*Ec* rpoZ as described (Hudson et al., 2009).

The *Ec* Crl was over-expressed in *E. coli* BL21(DE3) cells (Novo protein, Inc.) carrying pET28a-TEV-Crl. Protein expression was induced with 0.5 mM IPTG at 18 °C for 14h when OD_600_ reached to 0.6-0.8. Cell pellet was lysed in lysis buffer (50 mM Tris-HCl pH 7.7, 500 mM NaCl, 5% (v/v) glycerol, 5 mM β-mercaptoethanol, and protease inhibitor cocktail (Biomake.cn. Inc.)) using an Avestin EmulsiFlex-C3 cell disrupter (Avestin, Inc.). The lysate was centrifuged (16,000 g; 50 min; 4 °C) and the supernatant was loaded onto a 2 ml column packed with Ni-NTA agarose (SMART, Inc.). The proteins bound on resin were washed by the lysis buffer containing 20 mM imidazole and eluted with the lysis buffer containing 300 mM imidazole. The eluted fractions were mixed with TEV protease and dialyzed against 20 mM Tris-HCl, pH 7.7, 50 mM NaCl, 1mM DTT. The sample was reloaded onto a Ni-NTA column to remove his tag. The Crl was further purified on a Q HP column (HiPrep Q HP 16/10, GE healthcare Life Sciences) with a salt gradient of buffer A (20 mM Tris-HCl, pH 7.7, 50 mM NaCl, 1 mM DTT) and buffer B (20 mM Tris-HCl, pH 7.7, 1 M NaCl, 1 mM DTT). The fractions containing target proteins were collected, concentrated, and stored at −80 °C. The Crl and σ^S^ derivatives were prepared by the same procedure.

### Nucleic-acid scaffolds

The nucleic-acid scaffold for cryo-EM study was prepared by mixing synthetic nontemplate DNA, template DNA, and RNA at molar ratio of 1: 1.2: 1.5 and subjected to an annealing procedure (95°C, 5 min followed by 2°C-step cooling to 25°C) in annealing buffer (5 mM Tris-HCl, pH 8.0, 200 mM NaCl, and 10 mM MgCl_2_).

### Complex reconstitution of *E. coli* Crl-TAC

The Crl-σ^S^ binary complex was prepared by incubating Crl and σ^S^ at a molar ratio of 2:1 and purified by using a Superdex 75 gel filtration column (GE Healthcare). The *E. coli* Crl-TAC was assembled by directly incubating RNAP core enzyme, the Crl-σ^S^ binary complex, and the nucleic-acid scaffold at a molar ratio of 1:4:4 at 4 °C overnight. The mixture was loaded onto a Superdex 200 gel filtration column (GE Healthcare) and eluted with 10 mM HEPES pH 7.5, 50 mM KCl, 5 mM MgCl_2_, 3 mM DTT. Fractions containing *E. coli* Crl-TAC were collected and concentrated to ~12 mg/ml.

### Cryo-EM structure determination of *E. coli* Crl-TAC

The *E. coli* Crl-TAC sample was freshly prepared as described above and mixed with CHAPSO (Hampton Research, Inc.) to a final concentration 8 mM prior to grid preparation. About 4 μL of the complex sample was applied onto the glow-discharged C-flat CF-1.2/1.3 400 mesh holey carbon grids (Electron Microscopy Sciences) and the grid was plunge-frozen in liquid ethane using a Vitrobot Mark IV (FEI) with 95% chamber humidity at 10 °C.

The data were collected on a 300 keV Titan Krios (FEI) equipped with a K2 Summit direct electron detector (Gatan). A total of 3290 images of Crl-TAC were recorded using the Serial EM (Mastronarde, 2005) in super-resolution counting mode with a pixel size of 0.507 Å, and a dose rate of 6.7 electrons/pixel/s. Movies were recorded at 250 ms/frame for 8 s (32 frames total) and defocus range was varied between 2.0 μm and 2.5 μm. Frames in individual movies were aligned using MotionCor2 (Zheng et al., 2017), and Contrast-transfer-function estimations were performed using CTFFIND (Rohou and Grigorieff, 2015). About 1,160 particles were picked and subjected to 2D classification in RELION 3.0 (Fernandez-Leiro and Scheres, 2017). The resulting distinct two-dimensional classes were served as templates and a total of 315,977 particles for Crl-TAC were picked out. The resulting particles were manually inspected and subjected to 2D classification in RELION 3.0 by specifying 100 classes (Zivanov et al., 2018). Poorly-populated classes were removed. We used a 50-Å low-pass-filtered map calculated from structure of *E. coli* σ^S^ -TIC (Kang et al., 2017) (PDB: 5IPL) as the starting reference model for 3D classification. The final maps were obtained through 3D auto-refinement, CTF-refinement, Bayesian polishing, and post-processing in RELION 3.0 (Fig. S2). Gold-standard Fourier-shell-correlation analysis (FSC) (Henderson et al., 2012) indicated a mean map resolution of 3.80 Å for *Ec* Crl-TAC.

The crystal structure of *E. coli* σ^S^ -TIC (Kang et al., 2017)(PDB: 5IPL), crystal structure of *P. mirabilis* Crl (Cavaliere P et al., 2014) (PDB: 4Q11) were manually fit into the cryo-EM density map using Chimera (Pettersen et al., 2004b). Rigid body and real-space refinement was performed in Coot (Emsley and Cowtan, 2004) and Phenix (Adams et al., 2010).

### Hydrogen–deuterium exchange mass spectrometry (HDX-MS) of σ^S^

The HDX-MS was performed as recommended in (Masson et al., 2019). Amide hydrogen exchange of σ^S^ alone were started by diluting 3 μl protein sample at 19 μM into 27 μl D_2_O buffer (20 mM Tris, pH 7.7, 150 mM NaCl, 1mM TCEP) at 20 °C. At different time points (0 s, 10 s, 60 s, 300 s and 900s), the labeling reaction was quenched by the addition of chilled quench buffer (200 mM KH_2_PO4/K_2_HPO4, pH 2.2) and the reaction mixture was immediately frozen in liquid nitrogen. For the HDX-MS of σ^S^ in the presence of Crl, 50 μL σ^S^ at 34 μM were first mixed with 40 μl Crl at 176 μM and incubated at room temperature for 1 hr. 3 μL mixture was then labeled by adding 27 μL D_2_O buffer before being quenched at above time points and flash-frozen. All frozen samples were stored at −80 °C until analysis.

The thawed samples were immediately injected into HPLC-MS (Agilent 1100) system equipped with in-line peptic digestion and desalting. The desalted digests were then separated with a Hypersil Gold™ C18 analytical column (ThermoFisher) over a 19 min gradient and directly analyzed with an Orbitrap Fusion mass spectrometer (ThermoFisher). The HPLC system were extensively cleaned with blank injections between samples to minimize any carryover. Peptides identification was performed by tandem MS/MS under orbi/orbi mode. All peptides were identified using the Proteome Discoverer™ Software (ThermoFisher). We carried out the initial analysis of the peptide centroids with HD-Examiner v2.3 (Sierra Analytics) and then manually verified every peptide to check retention time, charge state, m/z range and the presence of overlapping peptides. The peptide coverage of σ^S^ were found to be 95.2% and the relative deuteration levels (%D) of each peptide were automatically calculated by HD-Examiner with the assumption that a fully deuterated sample retains 90% D in current LC setting.

### Yeast two-hybrid assay

The GAL4-based yeast two-hybrid system (MATECHMAKER GAL4 two-hybrid system3, Clontech Laboratories, Inc.) was used to analyze the protein-protein interaction according to the standard procedure. Briefly, wild-type or derivatives of *Ec*Crl and *Ec*σ^S^_2_ were cloned into prey vector pGADT7 and the bait vector pGBKT7, respectively. The bait and prey vectors were transformed into Y187 and AH109 yeast cells, respectively. The haploid colonies of Y187 were mated with haploid prey colonies of AH109 in the YPDA medium for 24h, and the diploid yeast cells containing both bait and prey vectors were selected on SD (–leu, –trp) plates at 30 °C for 48 h. The colonies were inoculated into SD (–Leu, –Trp) medium and cultured at 30 °C for 24 h. The resulting cell suspensions with a series of dilution were spotted onto SD (-Leu, -Trp) and SD (-Ade, -His, -Leu, -Trp) plates, incubated at 30 °C for 4-5 days. Positive colonies appear after 3 days on SD (-Ade, -His, -Leu, -Trp) plates and the plate images were taken on day 5-6 day after plating.

### Fluorescence labeling

The *E. coli* σ^S^(A239C) was labeled with fluorescein at residues C239. The labeling reaction mixture (2 mL) containing σ^S^ (0.07 mM) and Fluorescein-5-Maleimide (0.7 mM; Thermo Scientific, Inc.) in 10 mM Tris-HCl, pH 7.7, 100 mM NaCl, 1% glycerol was incubated overnight at 4℃, and the reaction was terminated by addition of 2 μL 1M DTT, and loaded onto a 5 mL PD-10 desalting column (Biorad, Inc.). The fractions containing labeled protein was pooled and concentrated to 3 mg/mL. The labeling efficiency is estimated ~ 70%.

### Fluorescence polarization

The reaction mixtures (100 μL) contain the fluorescein-labeled σ^S^ (4 nM; final concentration) with or without Crl (1 μM; final concentration) in FP buffer (10 mM Tris-HCl, pH 7.9, 300 mM NaCl,1 mM DTT, 1% glycerol, and 0.025% Tween-20) were incubated for 10 min at room temperature. RNAP core enzyme (0 nM to 1024 nM; final concentration) was added and incubated for 10 min at room temperature. The FP signals were measured using a plate reader (SPARK, TECAN Inc.) equipped with excitation filter of 485/20 nm and emission filter of 520/20 nm. The data were plotted in SigmaPlot (Systat software, Inc.) and the dissociation constant *K*d were estimated by fitting the data to the following equation,

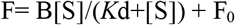

Where F is the FP signal at a given concentration of RNAP, F_0_ is the FP signal in the absence of RNAP, [S] is the concentration of RNAP, and B is an unconstrained constant.

### Stopped-flow assay

The stopped-flow assay was performed essentially as in (Feklistov et al., 2017). Briefly, to monitor the promoter melting by *Ec* σ^s^-RNAP holoenzymes in the presence or absence of Crl, 60 μL *Ec* σ^s^-RNAP holoenzyme (200 nM; final concentration) and 60 μL Cy3-λ P_R_ promoter DNA (20 nM; final concentration) in 10 mM Tris-HCl, pH 7.7, 300 mM NaCl, 10 mM MgCl_2_, 1 mM DTT were rapidly mixed and the change of Cy3 fluorescence was monitored in real time by a stopped-flow instrument (SX20, Applied Photophysics Ltd, UK) equipped with a excitation filter (515/9.3 nm) of and a long-pass emission filter (570 nm). Crl (1 μM; final concentration) was pre-incubated with *Ec* σ^s^-RNAP holoenzyme when indicated.

### *In vitro* transcription assay

The *osmY* promoter with tR2 terminator was prepared by PCR amplification of *osmY* promoter region (−107/+50 relative to the transcription start site) from *E. coli* genomic DNA using primers (forward primer: 5’- TTCCCTTCCTTATTAGCCGCTT -3’; reverse primer: 5’- AAATAAAAAGGCCTGCGATTACCAGCAGGCCCTGATATCTACGCATTGAACGG -3’). All of the *in vitro* transcription reactions were performed in transcription buffer (40 mM Tris-HCl, pH 7.9, 75 mM KCl, 5 mM MgCl_2_, 12.5%Glycerol, 2.5mM DTT) in a 20 μl reaction mixture.

To study the effect of overall transcription activation by Crl, Crl (500 nM; final concentration) was pre-incubated with σ^S^ (200 nM; final concentration) for 10 min at 30°C in transcription buffer prior to addition of RNAP core enzyme (100 nM; final concentration). Promoter DNA (100 nM; final concentration) was subsequently added and incubated for 10 min. RNA synthesis was allowed by addition of NTP mixture (30 μM ATP, 30 μM CTP, 30 μM GTP, 30 μM [α-^32^P]UTP (0.04 Bq/fmol); final concentration) for 15min at 30 °C. The reactions were terminated by adding 5 μL loading buffer (8 M urea, 20 mM EDTA, 0.025% xylene cyanol, and 0.025% bromophenol blue), boiled for 2 min, and cooled down in ice for 5 min.

To study the transcription activation of pre-assembled σ^s^-RNAP holoenzyme by Crl, reaction mixture containing pre-assembled *E. coli* σ^s^-RNAP (200 nM; final concentration) and Crl (0 nM to 1600 nM; final concentration) were incubated in transcription buffer for 10 min at 30 °C, then promoter DNA (200 nM; final concentration) was added and incubation for 10 min at 30 °C for open complex formation. RNA synthesis was allowed by the addition of NTP mixture (30 μM ATP, 30 μM CTP, 30 μM GTP, 30 μM [α-^32^P]UTP (0.04 Bq/fmol) for each; final concentration) for 15min at 30 °C. The reactions were terminated by adding 5 μL loading buffer (8 M urea, 20 mM EDTA, 0.025% xylene cyanol, and 0.025% bromophenol blue), boiled for 2 min, and cooled down in ice for 5 min. The RNA transcripts were separated by 15% urea-polyacrylamide slab gels (19:1 acrylamide/bisacrylamide) in 90 mM Tris-borate (pH 8.0) and 0.2 mM EDTA and analyzed by storage-phosphor scanning (Typhoon; GE Healthcare, Inc.).

### Quantification and statistical analysis

All biochemical assays were performed at least 3 times independently. Data were analyzed with SigmaPlot 10.0 (Systat Software Inc.).

